# First moment of lateral pressure in lipid membranes: theoretical calculation

**DOI:** 10.1101/2025.03.19.644141

**Authors:** Anna A. Drozdova

## Abstract

Analytical expression for the first moment of lateral pressure in a flat and bent membrane has been represented in this study. The calculation was carried out within the microscopic model of lipid bilayer. Macroscopic parameters of lipid membrane in this model (partial function, free energy, bending modulus, lateral pressure and so on) may be expressed through the microscopic parameters of the lipid chain (area per chain, length, bending rigidity of lipid). Lateral pressure profile in a lipid bilayer is a distribution of the entropic repulsion between neighboring lipid chains by membrane thickness. Lateral pressure in a bent bilayer is a very important characteristic of lipid membrane because it influences on membrane elastic constants. And although bending modulus is a well-known feature, little is known about the position of neutral surface (non-expanded surface). Moreover, analytical expression for lateral pressure in the bent bilayer is connected to the expression of mixed deformation modulus (bending with stretching). Its zero value indicates the case of pure bending and relaxation of neutral surface. In contradistinction to membrane with curvature the first moment of lateral pressure in a flat membrane (so named spontaneous bending moment) is significantly affected on membrane protein activity because it is related with macroscopic characteristic – inter-facial surface tension. So, the microscopic theory represented in this article enables calculation of bending modulus for lipid bilayer, bending modulus for lipid monolayer, area stretching modulus, bending-stretching modulus, spontaneous bending moment for lipid monolayer and also defining the position of neutral surface.

## I. Introduction

The relevance of the present study is supported by the fact that biological membranes surround all living cells and form them in closed organelles. The basic structural element of membrane is a lipid bilayer. Its mechanical and thermodynamic properties are supposed to play a significant role in the functioning of organized matter.

Lipids typically have one polar (hydrophilic) “head” and two non-polar (hydrophobic) “tails” (hydrocarbon chains). There is a great variety of membrane constituents and lipid characteristics (head group type, chain length *etc*). Theoretical study of thermodynamic characteristic for lipid membrane has been represented in this article. Analytical theory for lipid membranes is based on the microscopic model that was earlier developed by colleagues [1–3]. According to this theory each lipid chain is represented as a flexible string of finite thickness (incompressible area per chain) with fixed bending rigidity (the case of persistence length comparable with the chain full length is considered). The ensemble of lipid chains is investigated within a mean-field approximation with respect to string collisions. As per this approach an entropic repulsion between lipid tails is represented as an effective lateral potential acting on each string in each monolayer. The negative lateral tension keeping all the strings (i.e. lipids) together comes from hydrophilic-hydrophobic interface in the head-group area of a membrane and is treated as an input parameter of the model. Thus the energy functional for a lipid chain can be written analytically and then partition function and free energy of the chain can be calculated. The analytical expression of the free energy enables to obtain any macroscopic characteristics of lipid bilayer, for example, the lateral pressure profile that proves to be a very important feature of lipid membrane because it is related to the elastic properties of lipid bilayer [4] and also the functionality of mechanosensitive ion channels [5–7].

Elasticity of lipid membrane will be described in this article. One can write the density of bending energy in Helfrich form [8]:

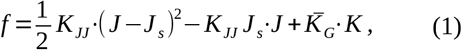

where *J* is a mean curvature, *J* _*s*_ is a spontaneous curvature, *K* is a Gaussian curvature. The two constants that control the membrane shape are bending modulus *K*_*JJ*_ and saddle-splay modulus 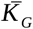 (index “*JJ*” means a second derivative of the free energy with respect to curvature 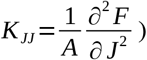. In this research author does not consider the topology of membrane. So, the last term in eq. (1) will be omitted.

It is important to note that Eq. (1) represents a pure bending of lipid membrane. The bending of lipid bilayer usually accompanies by the stretching. However, it is possible to calculate independently area stretching deformation 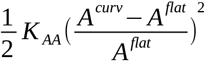 and bending deformation 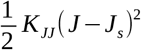 without mixed deformation 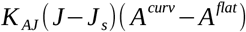. In such case the neutral surface has to be relaxed [9–11] (The neutral surface is defined as surface of zero lateral expansion under bending deformation of the membrane).

Experimental and numerical calculations of the bending modulus are unable to determine the exact position of the neutral surface. However, it was established within the experimental studies [9] that it is possible to calculate the energy of pure bending only by minimizing of the lipid energy in a curved monolayer with respect to the position of the neutral surface. Generally, this fact complicates the calculation by the formula: 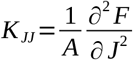. But calculation of the bending modulus through the first moment of lateral pressure enables to solve this problem, because bending modulus *K*_*JJ*_ is connected to lateral pressure through moment of lateral pressure [4, 12].

In addition, theoretical calculation of the pressure profile enables to calculate the modulus of mixed deformation *K* _*AJ*_. The point is that *K* _*AJ*_ is directly related to the position of the neutral surface [10]. And displacement of neutral surface will be discussed in this article within several models of bending (various models of bending differ by the position of the neutral surface).

## II. Methods

### 1. Flexible string model

Microscopic model of lipid molecule as an ensemble of flexible strings was investigated by colleagues in work [1–3]. This theory enables to express any macroscopic characteristics of membrane (free energy, lateral pressure, surface tension and so on) through microscopic parameters of lipid molecule (area per lipid chain, monolayer thickness). Some aspects of the theory will be briefly described in this section.

Hydrocarbon lipid chain is considered as Euler’s elastic beam of finite thickness, and its energy functional is written in the following form:

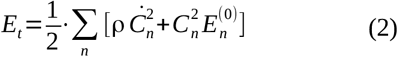

where ρ is the linear density of the lipid chain, 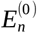 and *C*_*n*_ are, respectively, eigen values and amplitudes of eigen functions 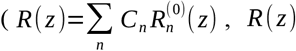 is the deviation of the beam from the straight line at each level *z* - see fig. 1 in [2]) of the conformation energy operator 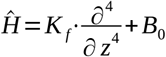. Here the first term is the bending energy of the chain and the second term is the coefficient of entropic repulsion with the neighboring chains. *K*_*f*_ is the bending (flexural) rigidity of lipid chain. Its estimate was taken is the following form: *K*_*f*_ ≈*k* _*B*_ *T*·*l*_*p*_, where persistence length - *l* _*p*_=*L*_0_ for short, *l* _*p*_=*L*_0_ / 2 for long and *l* _*p*_=*L*_0_ / 3 for very long hydrocarbon lipid chains [1, 13–16], and *L*_0_ is a monolayer thickness.

**Figure 1.**
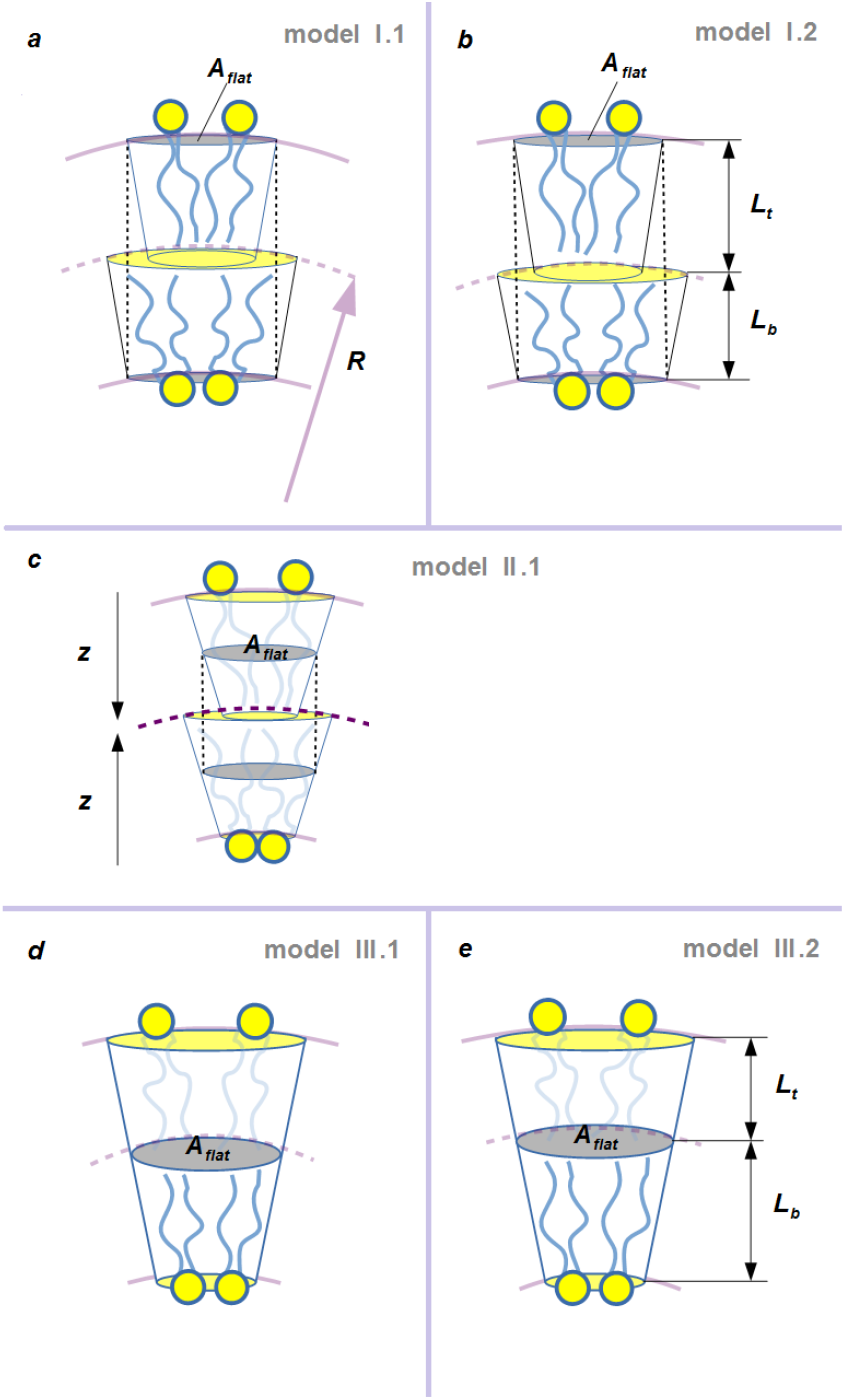
Bending model of lipid bilayer. Here *R* is a curvature radius and 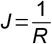 is a curvature of inter-monolayer surface (dotted line at all sketches). *L*_*b*_ is a thickness of the inner (‘bottom’) monolayer, *L*_*t*_ is a thickness of the outer (‘top’) monolayer, *L*_*b*_+*L*_*t*_=2*L*_0_. At all sketches *A*_*flat*_≡ *A* is area per lipid chain in the flat bilayer – see Eqs. (17), (19), (20) and zero of axis *z* begins at the hydrophilic-hydrophobic interface. *a, b* – neutral surfaces are located in hydrophilic-hydrophobic region; *c* – neutral surfaces are located in the middle of each monolayer (this is a case of pure bending); *d, e* – neutral surface is located in the inter-monolayer surface. For bending models that are represented in sketches *b* and *e* condition of the volume incompressibility leads to difference in thicknesses of monolayers *L*_*b*_≠*L*_*t*_. For other models of bending *a, c* and *d* thicknesses of the bot and top monolayers are assumed to be equivalent *L*_*b*_=*L*_*t*_ During deformations in sketches *a* and *d* neutral surfaces are being shifted along to the monolayer depth (fig. 2).

Knowing the energy functional (2) one can calculate the partition function consisted of the product of two equal components, 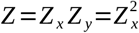, where *Z*_*x*_ in the amplitude representation {*C*_*n*_} takes the following form:

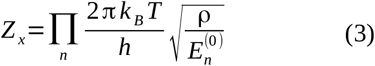

Next, one can write the free energy of lipid tails:

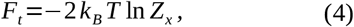

Substituting Eq. (3) into (4) and doing some transformations one can obtain:

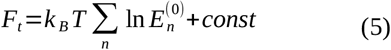

Next, the lateral pressure inside the hydrophobic part of the bilayer can be calculated by differentiating of the free energy with respect to area per chain 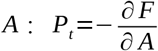 and the lateral pressure profile 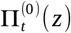 in a flat bilayer is introduced in the following form:

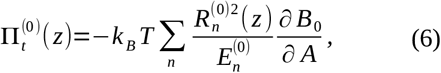

where entropic repulsion coefficient in a flat membrane *B*_0_ and the area per lipid chain *A* can be found from the self-consistency equation [2]:

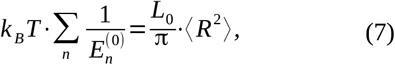

where 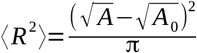 is the average of square deviation. Using relation between incompressible area per lipid chain *A*_0_ =0,2 *nm*^2^ and is the real area occupied by the chain *A* : *A*=*a*·*A*_0_ one can write Eq. (7) in dimensionless form:

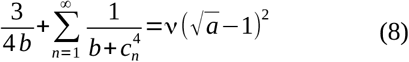

Here 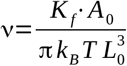 and 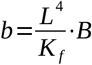 are dimensionless coefficients, *c*_*n*_=π *n*−π/ 4 [2]. Also in Eq. (6) *z* is the distance (depth) from head-group region inside lipid monolayer.

The self-assembly condition (*i*.*e*. zero total lateral tension of the self-assembled membrane) is represented in the form:

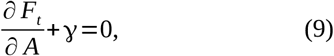

where γ is the surface tension at the interface between the polar and hydrophobic regions of the membrane. The physical meaning of this value is the surface tension at the lipid-water interface (situated between hydrophobic tails and hydrophilic heads): γ=30−100 *mN* /*m*. The equilibrium condition (9) simply states that the repulsion between chains should be balanced by the surface tension γ :

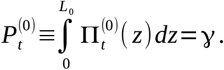

For calculation of thermodynamic characteristics in a bent lipid membrane one should obtain the analytical expression for the correction of lateral pressure profile due to the curvature *J* [17]. Free energy of the bent membrane is expressed by Equation:

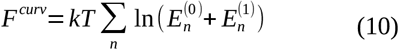

Here 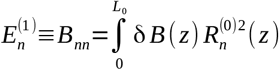 is the first correction to eigen value 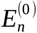 [18], where 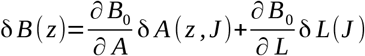 perturbation of self-consistent coefficient due to the curvature (it is assumed *L*_0_·*J* ≪1): δ *A* (*z, J*) is z-dependent area change caused by the membrane bending and δ *L*(*J*) is a change of the monolayer thickness.

Hence, differentiating Eq. (10) one can obtain the first correction to the lateral pressure profile in bent membrane with finite curvature

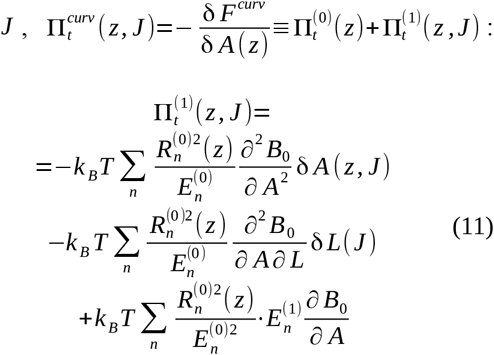

It is important to note that integration of the first correction (11) results in the lateral pressure in curved monolayer 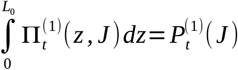.

### 2. The first moment of lateral pressure profile

At the beginning one need define the deformation moduli [4, 10]:

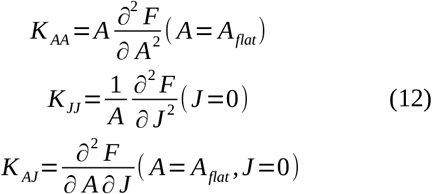

Here *A*_*flat*_ is the area per chain in the flat membrane, *K* _*AA*_ is the area compressibility modulus, *K* _*JJ*_ - bending modulus and *K* _*AJ*_ - modulus of mixed deformation.

According to the studies of D. Marsh and W. Helfrich [4, 12] the bending modulus of the monolayer is related to the first moment of the lateral pressure profile in a curved monolayer:

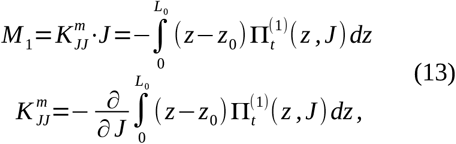

where *z*_0_ is the position of the neutral surface.

Analogously, the spontaneous bending moment is given in the following form [12]:

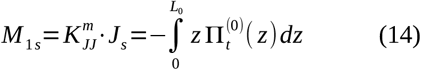

Here *J* _*s*_ is the spontaneous curvature. Its physical meaning arises from geometrical difference in size of hydrocarbon chain and polar head. With addition of some non-bilayer lipids the bilayer acquires a non-zero spontaneous curvature. Also a spontaneous curvature may be produced by changing in pH [12]. It is important to note that spontaneous bending moment does not depend on the location of the neutral surface because its magnitude for inner (‘bottom’) and outer (‘top’) monolayers of the membrane is differ by sign. In addition there is a difference in signs for the corrections to the lateral pressure 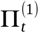, but there is no such difference in signs for the bending moment for the inner and outer monolayers *M* _1_. Hence the position of the neutral surface will be kept in mind during the calculation of the first bending moment *M* _1_ (13).

The mixed deformation modulus *K* _*AJ*_ can be calculated analytically within the flexible string model. The general idea of this calculation arises from expression for the free energy of bent membrane:

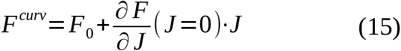

Differentiating of Eq. (15) with respect to area per chain *A* results in the relation between the first correction to the lateral pressure and the modulus of mixed deformation (*cf*. with Eq. (12)):

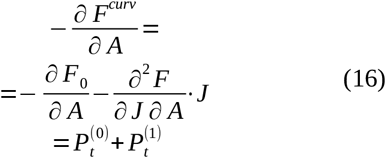

### 3. Bending model

In this section the model of bent bilayer will be introduced. This can be done by considering area per lipid being dependent on *z*: *A*^*curv*^= *A*+δ *A* (*z*).

Figure 1 represents a bending model for coupled and uncoupled monolayers with different location of the neutral surface with regard to the middle of bilayer. If neutral surfaces are located in the hydrophilic-hydrophobic interfaces of the membrane [19] then area per lipid chain is represented by expression [20]:

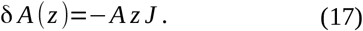

Here *A* is the area per chain in a flat bilayer. Further, solving equation of the lipid volume conservation [2, 19] one can obtain the thickness *L* of curved monolayer, *L*=*L*_0_+δ *L*:

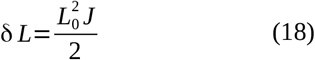

Fig. 1 a illustrates bending of a bilayer described by Eq. (17), and Fig. 1 b illustrates bending of a bilayer with conservation of volume per lipid chain - Eqs. (17) and (18).

If neutral surfaces are located in the middle of the monolayer [21] then area per lipid chain is represented by expression:

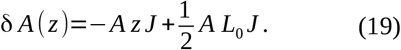

Using condition for the conservation of volume per lipid molecule one can obtain that δ *L*=0 up to a first order in curvature (see fig. 1 c).

If neutral surfaces are located in the middle of bilayer [12, 22] then area per lipid chain is represented by expression:

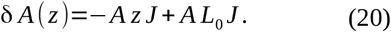

Using conservation of volume per lipid chain one can obtain the thickness *L* of the curved monolayer, *L*=*L*_0_+δ *L*:

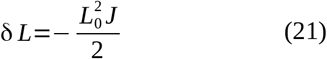

Relation of Eqs. (17) - (21) hold for both monolayers provided that opposite signs are assigned to the mean curvatures of the opposite monolayers of the curved bilayer.

Fig. 1 d illustrates bending of a bilayer described by Eq. (20), and Fig. 1 e illustrates bending of a bilayer with conservation of the volume per lipid chain - Eqs. (20) and (21).

### 4. Location of the neutral surface

The microscopic theory of flexible strings enables to define the position of neutral surface. According to Eqs. (12) and (16) the modulus of mixed deformation can be written in the following form:

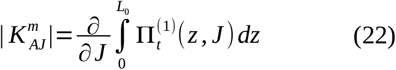

If its value is zero then the neutral surface is relaxed 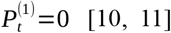. Fig. 1 c illustrates this case (pure bending). For other cases the neutral surface has been shifted along to the monolayer depth. (for other cases *K* _*AJ*_ ≠0) This displacement of the neutral surface can be derived from calculated lateral pressure profile (see fig. 2 a-e). Obviously, that the point in which the correction to lateral pressure is zero indicates the position of neutral surface.

**Figure 2.**
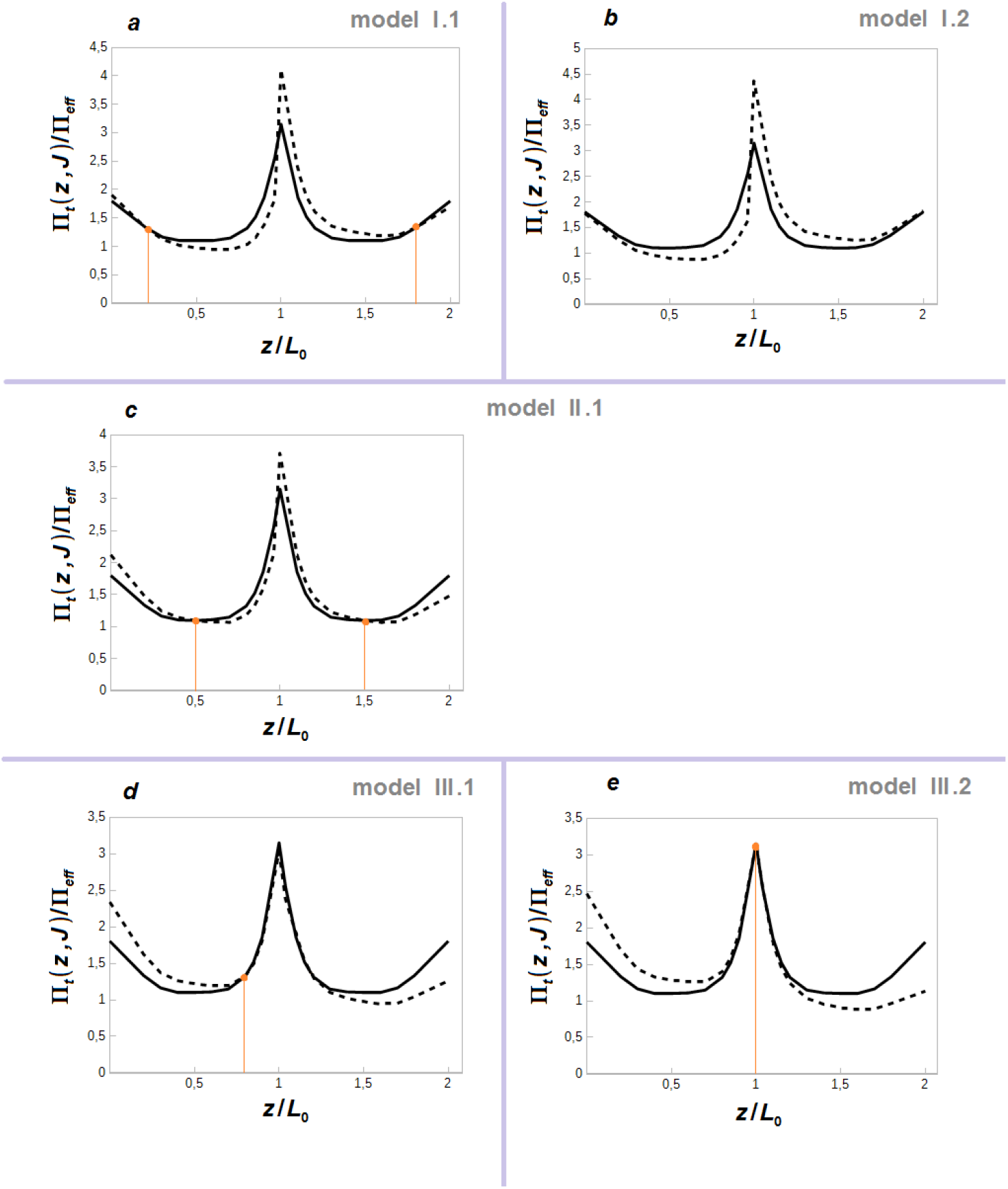
Lateral pressure profile in hydrophobic part of lipid bilayer normalized by Π_*eff*_=γ/ *L*_0_. Solid line – Π_0_(*z*) - lateral pressure profile in a flat bilayer; dotted line – Π(*z, J*) - lateral pressure profile in a bilayer with curvature *J, R*=20 *nm*. γ is inter-facial surface tension, γ=0,05 *N* /*m*. *a* – lateral pressure in lipid membranes according to bending model I.1; *b* – lateral pressure profile in lipid membrane according to bending model I.2 and other sketches (*c, d, e*) may be described similarly. It is important to note that the points, where curves Π(*z, J*) and Π_0_(*z*) coincides with each other, defines location of the neutral Surfaces 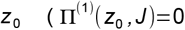, there are vertical lines at all sketches). The position of neutral surface is altered in different bending models. For bending model I.1 *z* _0_/ *L*_0_=1 /5, but for bending model III.1 *z* _0_/ *L*_0_=4/ 5. For other models in which a volume per molecule is incompressible the position *z* _0_ coincides with initial location before deformation: *z* _0_/ *L*_0_=0 for I.2 model; *z* _0_/ *L*_0_=1 / 2 for II.1 model; *z* _0_/ *L*_0_=1 for III.2 model.

## III. Results

The first moment of lateral pressure profile – see Eqs. (13)–(14) – will be calculated in this section.

Substituting Eq. (6) into (14) one can calculate the spontaneous bending moment for inner (‘bottom’) monolayer:

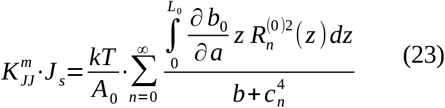

Integration and summation in Eq. (23) may be interchanged since it is carried out with respect to independent variables. Computing integral in Eq. (23) one gets 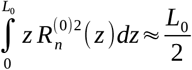 for large *n*. So, analytical expression for spontaneous bending moment is given by Eq.:

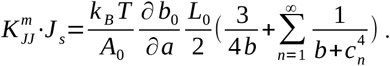

Accounting for Eqs. (6) and (9) one may obtain a more compact formula:

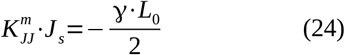

Table 1 represents calculated spontaneous bending moment for various lipids and different magnitudes of inter-facial surface tension γ. The obtained results are in a good agreement with molecular dynamic simulations by S. Ollila [23], where its value

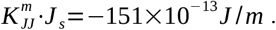

**Table 1.**
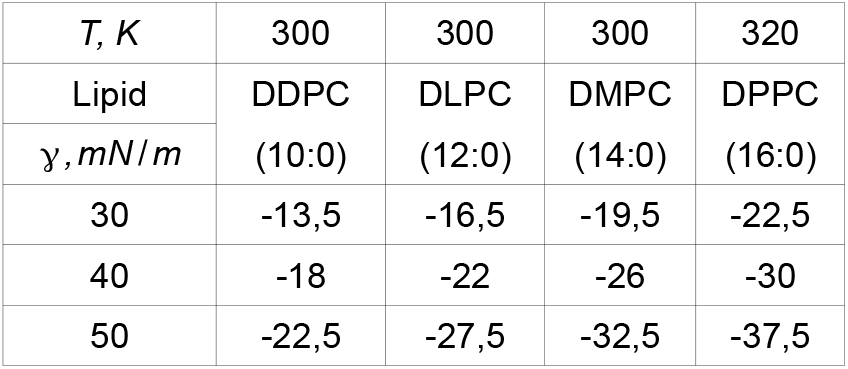
Spontaneous bending moment for inner (‘bottom’) monolayer 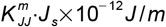 calculated analytically within a microscopic model of flexible strings. Incompressible area per chain *A*_0_ =0,2 *nm*^2^, a temperature *T* and inter-facial surface tension γ are input parameters for calculations. Here *L*_0_=0, 9 *nm* - a hydrophobic thickness of monolayer in DDPC (10:0) lipids, *L*_0_=1, 1 *nm* - a hydrophobic thickness of monolayer in DLPC (12:0) lipids, *L*_0_=1, 3 *nm* - a hydrophobic thickness of monolayer in DMPC (14:0) lipids and *L*_0_=1, 5 *nm* - a hydrophobic thickness of monolayer in DPPC (16:0) lipids.

| $T, K$ | 300 | 300 | 300 | 320 |
| --- | --- | --- | --- | --- |
| Lipid | DDPC | DLPC | DMPC | DPPC |
| $\gamma, mN/m$ | (10:0) | (12:0) | (14:0) | (16:0) |
| 30 | -13,5 | -16,5 | -19,5 | -22,5 |
| 40 | -18 | -22 | -26 | -30 |
| 50 | -22,5 | -27,5 | -32,5 | -37,5 |

Next, substituting expression for the correction to lateral pressure profile (11) into Eq. (13) and calculating integral 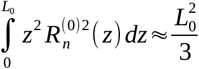 (see Supplementary text in [24]) one gets a bending modulus in different bending model (*i*.*e*. Eqs. (17) - (21) and fig. 2 a-e). Calculated results for various fully saturated lipids are represented in Table 2. Usually experimental value for bending modulus in different lipid bilayers vary between (0, 1−6)×10^−19^ *J* [25–27]. So, calculation within flexible string model gives a good agreement with these data.

**Table 2.** Bending modulus of lipid bilayer calculated analytically in different bending models (e.g. Eqs. (17) - (21) and fig. 2)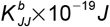. Models I.1 and III.1 represent a bending of monolayers when neutral surfaces are non-relaxed and shifted against the original position. The model II.1 represents a pure bending of uncoupled monolayers. Models I.2 and III.2 represents a bending with additional condition of the volume conservation. Incompressible area per chain *A* =0,2 *nm*^2^, a temperature *T* and inter-facial surface tension γ are input data for the calculations. Here *L*_0_=0, 9 *nm* - a hydrophobic thickness of monolayer in DDPC (10:0) lipids, *L*_0_=1, 1 *nm* - a hydrophobic thickness of monolayer in DLPC (12:0) lipids and so on.

| $T, K$ | 300 | 300 | 300 | 320 | 333 | 343 | 350 |
| --- | --- | --- | --- | --- | --- | --- | --- |
| Lipid | DDPC | DLPC | DMPC | DPPC | DSPC | DAPC | DBPC |
| $\gamma, mN/m$ | (10:0) | (12:0) | (14:0) | (16:0) | (18:0) | (20:0) | (22:0) |
| Bending Model I.1, III.1 |  |  |  |  |  |  |  |
| 30 | 0,32 | 0,482 | 0,68 | 0,89 | 1,131 | 1,393 | 1,65 |
| 40 | 0,474 | 0,716 | 1 | 1,29 | 1,615 | 1,98 | 2,33 |
| 50 | 0,651 | 0,977 | 1,36 | 1,7 | 2,132 | 2,607 | 3,05 |
| Bending model II.1 |  |  |  |  |  |  |  |
| 30 | 0,17 | 0,26 | 0,36 | 0,48 | 0,6 | 0,74 | 0,87 |
| 40 | 0,26 | 0,39 | 0,54 | 0,69 | 0,86 | 1,05 | 1,23 |
| 50 | 0,35 | 0,53 | 0,73 | 0,91 | 1,14 | 1,39 | 1,62 |
| Bending Model I.2, III.2 |  |  |  |  |  |  |  |
| 30 | 0,5 | 0,754 | 1,07 | 1,42 | 1,81 | 2,24 | 2,68 |
| 40 | 0,73 | 1,106 | 1,56 | 2 | 2,56 | 3,15 | 3,74 |
| 50 | 0,99 | 1,494 | 2,09 | 2,66 | 3,35 | 4,11 | 4,87 |

It is important to note that DMPC (14:0) undergoes lipid gel phase transition at temperature *T* =296 *K*. So lipids with short hydrocarbon chains e.g. DDPC (10:0), DLPC (12:0) and DMPC (14:0) should be calculated at *T=*300 *K*. And lipids with long hydrocarbon chains e.g. DPPC (16:0), DSPC (18:0), DAPC (20:0) and DBPC (22:0) should be calculated at different temperatures (see Table 2).

There is the relation between bending *K*_*JJ*_ and stretching moduli *K*_*AA*_ of lipid bilayer that is expressed by following Eq. [2, 9]:

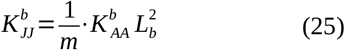

Obviously, that ratio *m* is different for various models of bending. For a start one need calculate area stretching modulus in flexible string model 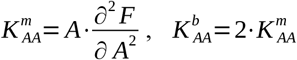. see Table 3. Magnitude of the area stretching modulus 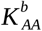 weakly depends on thickness of the bilayer and arises majorly from inter-facial surface tension γ. Hence, it has got ratio *m*=12 for bending models’ I.1 and III.1, *m*=24 – for bending model II.1 and *m*=8 – for bending models’ I.2 and III.2 (see fig. 3). It is obvious that bending model II.1 gives an agreement with polymer brush theory [9]. And in this section only expression of bending modulus for the case of pure bending (within bending model II.1) will be represented. Deformation consisted of bending and stretching will be go over in App. 1.

**Table 3.** Area compressibility modulus of bilayer 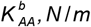. Analytical expression within a flexible string model is represented by the formula: 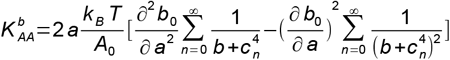 (see Eqs. (10), (12)). Here *L*_0_=0, 9 *nm* - a hydrophobic thickness of monolayer in DDPC (10:0) lipids, *L*_0_=1, 1 *nm* - a hydrophobic thickness of monolayer in DLPC (12:0) lipids and so on.

| T, K | 300 | 300 | 300 | 320 | 333 | 343 | 350 |
| --- | --- | --- | --- | --- | --- | --- | --- |
| Lipid | DDPC | DLPC | DMPC | DPPC | DSPC | DAPC | DBPC |
| $\gamma, mN/m$ | (10:0) | (12:0) | (14:0) | (16:0) | (18:0) | (20:0) | (22:0) |
| 30 | 0,122 | 0,123 | 0,125 | 0,124 | 0,122 | 0,121 | 0,118 |
| 40 | 0,179 | 0,182 | 0,183 | 0,178 | 0,174 | 0,171 | 0,166 |
| 50 | 0,245 | 0,247 | 0,247 | 0,235 | 0,229 | 0,225 | 0,217 |

**Figure 3.**
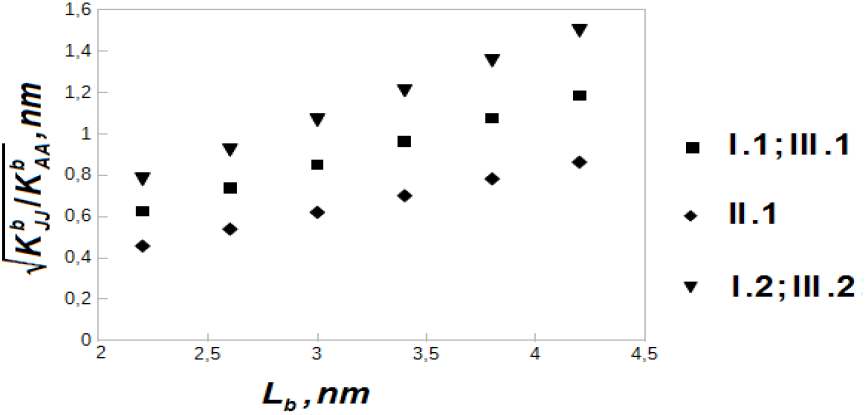
Dependence of membrane bending – to stretch modulus ratio on the hydrophobic thickness of fully saturated lipids at temperatures *T*=300 K for DLPC (12:0) and DMPC (14:0); *T*=320 K for DPPC (16:0); *T*=333 K for DSPC (18:0); *T*=343 K for DAPC (20:0); *T*=350 K for DBPC (22:0).

So, according to Eqs.(11), (13), (19) and (22) calculation of bending modulus for a pure bent monolayer would be simplified by condition of zero modulus of mixed deformation 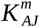:

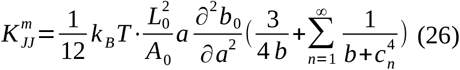

Here parameters of lipid molecules *a, b* and 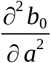 should be calculated from the self-consistency equation (8) (*viz*. App. 2). Bending modulus in Eq. (26) may be expressed through inter-facial surface tension

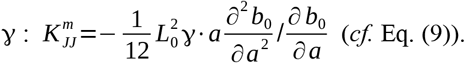

It is important to note that bending modulus for bilayer is always twice of bending modulus for monolayer 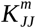 expressed by Eq. (13):

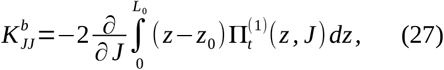

but not expression 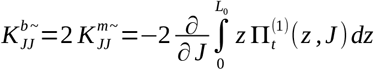 or equivalent formula represented in Eq. (12) without displacement of the neutral surface (see App. 1). This fact provides a well-known statement that bending modulus of bilayer is twice that of one of the monolayers if two monolayers are assumed to be uncoupled.

## IV. Discussion

Generally authors of many studies select the location of neutral surface *z*_0_ in different ways or state its location to be not fixed. In this research various models of membrane bending were considered. Those models differ by the location of neutral surface. It was assumed for the modeling of bilayer bending that area per lipid chain has acquired *z*-dependent component. Then variation of the area per chain was applied to get some macroscopic parameters for the bent lipid membrane. Namely, calculated analytically lateral pressure profile [17] is connected to the bending modulus through Eq. (13), (14). Moreover, theoretical calculation of lateral pressure profile resolves the definition of the neutral surface position and then calculation of mixed deformation modulus according to Eq. (22). [10]. These parameters are especially important in the estimation of relation between bending and stretching moduli 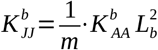, where ratio *m* = 24 for the case of pure bending [9]. At the same time calculations were performed for short- and long-tails fully saturated lipids and two monolayers are confirmed to be uncoupled. For coupled monolayers similar relation gives ratio *m*=12 [27]. Bending of the monolayers with neutral surfaces located in the hydrophilic-hydrophobic interface gives ratio *m*=12 [27]. Also it was shown, that spontaneous bending moment for lipid monolayer 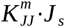 arises majorly from inter-facial surface tension and the monolayer thickness. This fact is well consistent with calculation in work [28].

## V. Conclusion

The microscopic theory of lipid bilayer with finite curvature was first developed in this study.

Free energy of lipid membrane was calculated analytically within this theory and then other relevant features were estimated *viz*. lateral pressure, bending modulus for the monolayer and bending modulus for the bilayer, and also spontaneous bending moment.

Since analytical formula for the first bending moment describes the bending of membrane it can be helpful in a getting of energy difference between various conformation states of membrane protein. In comparison with the curved membrane the first moment of lateral pressure in the flat membrane strongly influences on the gating mechanism of mechanosensitive channels and other proteins. This is caused by the fact that large hydrophobic energy arisen from spontaneous bending moment will reduce opening energy barrier in the channel [24, 29–30].

Significance of the present research is also supported by the fact that experimental study of lateral pressure profile are difficult. And moments of lateral pressure are connected with membrane elastic constants. So, analytical expression for the moments of lateral pressure can provide a calculation of bending modulus for both monolayer and bilayer of the membrane within various bending models (that differ by the location of neutral surface). At last, accurate position of neutral surface can be estimated from curves of lateral pressure. The fact is that during the bending process neutral surface may be deformed (i.e. changed its location with respect to initial-one that was fixed at consideration of the bending model). In such cases the mixed deformation (bending with stretching) will be occur.

## Appendices

### Appendix 1

In this section analytical formula for bending modulus in various bending models will be obtained. Substituting Eqs. (17) in (11) and then obtained expression in 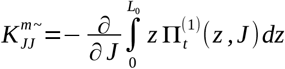 and (22) a reader obtains expression for bending and mixed deformation moduli: 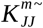 and 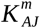.

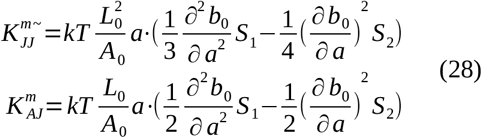

Here 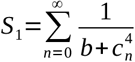 and 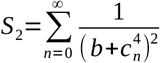 are sums over *n*, may be calculated with theory of residues: 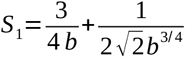 and Eq. (28) represents a bending model in Fig. 1 a and Fig. 2 a Combination 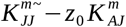 gives 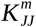 according to Eq. (13). At the same time 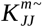 is equivalent to the second derivative of the free energy with respect to *J* – Eq. (12):

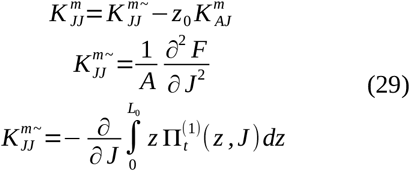

Analogously, similar expressions may be obtained for bending model I.2 that was represented in Fig. 1 b and Fig. 2 b. The difference is that 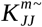 and 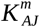 acquires components multiplied by derivatives of entropic potential with respect to thickness *L:* 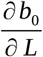 and 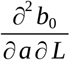.

Substituting Eqs. (20) in (11) and then obtained expression in 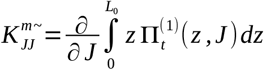 and (22) a reader obtains expression for bending and mixed deformation moduli 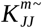 and 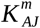 for bending model III.1 that was represented in Fig. 1 d and Fig. 2 d:

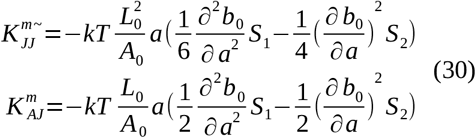

Model III.2 can be calculated similarly.

It is important to note that bending modulus by Eq. (30) (according to Eq. (11), *cf*. Eq. (12)) has obtained to be negative magnitude. This is caused by the fact that for model III.1 a relaxation of lipid asymmetry is occurred: 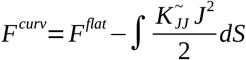. But bending modulus according to Eq. (13) has got to be positive value. If bending modulus of bilayer is represented by expression 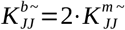, the ratio *m* in Eq. (25) takes value *m*=48 in model III.1. Similar relation in model III.2 gives ratio *m*=24.

### Appendix 2

In this section self-consistency parameters of lipid chain will be figured out. Area per lipid chain *a* and entropic potential coefficient *b* can be calculated from self-consistency equation (8) and equilibrium equation (9). Calculated result for area per lipid chain is represented in table 4. Entropic potential *b* can be expressed by the formula [1–2]:

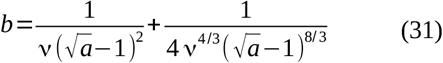

**Table 4.** Area per lipid chain *a* calculated analytically within microscopic model of flexible strings by using self-consistency equation (8) and equilibrium equation (9). Here lipids DDPC, DLPC, DMPC were calculated at flexural rigidity 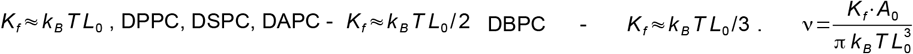 is dimensionless parameter, γ - inter-facial surface tension.

| Lipid | T, K | $\nu$ | $\gamma, mN/m$ | | |
| --- | --- | --- | --- | --- | --- |
|  |  |  | 30 | 40 | 50 |
| DDPC (10:0) | 300 | 0,0786 | 3,22 | 2,78 | 2,49 |
| DLPC (12:0) | 300 | 0,0526 | 3,29 | 2,84 | 2,55 |
| DMPC (14:0) | 300 | 0,0377 | 3,36 | 2,89 | 2,61 |
| DPPC (16:0) | 320 | 0,0142 | 3,74 | 3,23 | 2,91 |
| DSPC (18:0) | 333 | 0,011 | 3,91 | 3,38 | 3,04 |
| DAPC (20:0) | 343 | 0,0088 | 4,06 | 3,51 | 3,16 |
| DBPC (22:0) | 350 | 0,0048 | 4,4 | 3,79 | 3,4 |

Here dimensionless parameter ν one can see also in table 4.

At last, a reader can obtain expressions for derivatives of entropic coefficient required in expression for lateral pressure profile and bending modulus.

A differentiation of Eq. (8): 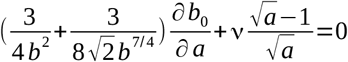 and following re-differentiation of obtained expression results in formula for the first and second derivatives of entropic potential 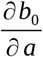 and 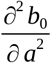:

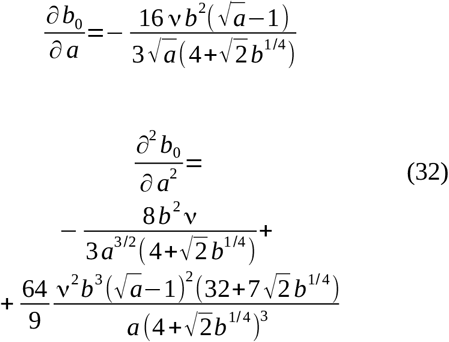

Analogously, mixed derivatives may be calculated from self-consistency equation (8):

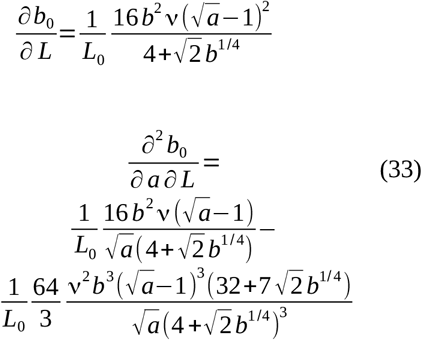

## Acknowledgments

This research did not receive any specific grant from funding agencies in the public, commercial or not-for-profit sectors.

## References

1. S. Mukhin and S. Baoukina, Phys Rev E, 71, 6 (2005)

2. Boris Kheyfets, Timur Galimzyanov, Anna Drozdova and Sergei Mukhin, Phys Rev E, 94, 042415 (2016)

3. S. I. Mukhin and B. B. Kheyfets, Phys Rev E, 82, 051901 (2010)

4. W. Helfrich, in Physics of Defects, North-Holland Publishing Company, Amsterdam, 716 (1981)

5. J. Gullngsrud and K. Schulten, Biophys J, 86, 3496 (2004)

6. Grischa R. Meyer, Justin Gullingsrud, Klaus Schulten and Boris Martinac, Biophys J, 91, 1630 (2006)

7. E. Perozo and D. C. Ress, Curr Opin Struc Biol, 13, 432 (2003)

8. W. Helfrich, Naturforsch C, 28 (11), 693 (1973)

9. W. Rawicz, K. C. Olbrich, T. McIntosh, D. Needham and E. Evans, Biophys J, 79, 328 (2000)

10. M. M. Kozlov, M. Winterhalter, J Phys II France, 1, 1085 (1991)

11. M. M. Kozlov, M. Winterhalter, Biol. Membr. 10, 71 (1993)

12. D. Marsh, Biophys J, 93, 3884 (2007)

13. P. J. Flory, J Chem Phys, 17, 303 (1949)

14. Polymer Encyclopedia (BSE, Moscow, 1997), vols. 1-3

15. P. Gittes, B. Mickey, J. Nettleton and J. Howard, J Cell Biol, 120, 923 (1993)

16. D. Nelson, Defects and Geometry in Condensed Matter Physics (Cambridge University Press, Cambridge, UK), 2002

17. A. A. Drozdova and S. I. Mukhin, JETP, accepted (2017)

18. L. D. Landau, E. M. Lifshitz. Quantum Mechanics (Pergamon, Oxford, 1970)

19. M. Hamm, M. M. Kozlov, Eur Phys J E, 3, 323 (2000)

20. Samuel A. Safran, Statistical Thermodynamics of Surfaces, Interfaces and Membranes, Addison-Wesley, Massachusetts, 1994

21. L. D. Landau and E. M. Lifshits, Course of Theoretical Physics, Theory of Elasticity, Vol. 7 (Butterworth-Heinemann, London), 2008

22. D. P. Siegel, M. M. Kozlov, Biophys J, 87(1), 366 (2004)

23. O. H. S. Ollila, H. J. Risselada, M. Louhivuori, E. Lindahl, I. Vattulainen and S. J. Marrink, Phys Rev Lett, 102, 078101 (2009)

24. Anna Drozdova, Sergei Mukhin, biorxiv preprint, doi: 10.1101/062463

25. Lee C. H., Lin W. C., Wang J., Phys Rev E, 64, 020901 (2001)

26. Seto H., Yamada N. L., Nagao M., Hishida M., Takeda M. Eur Phys J E Soft Matter, 26, 217 (2008)

27. D. Marsh, Chem Phys Lipids, 144, 146 (2006)

28. Nabina Mukherjee, MacDonald Jose, Jan Peter Birkner, Martin Walko, Helgi I. Ingolfsson, Anna Dimitrova, Clement Arnarez, Siewert J. Marrink and Armagan Kocer, FASEB J, 28 (10), 4992 (2014)

29. O. H. S. Ollila, M. Louhivuori, S. J. Marrink and I. Vattulainen, Biophys J, 100, 1651 (2011)

30. Anna A. Drozdova and Sergei I. Mukhin, Biophys J, 104(2), 244a (2013)

